# Tracking key virulence loci encoding aerobactin and salmochelin siderophore synthesis in *Klebsiella pneumoniae*

**DOI:** 10.1101/376236

**Authors:** Margaret M. C. Lam, Kelly L. Wyres, Louise M. Judd, Ryan R. Wick, Adam Jenney, Sylvain Brisse, Kathryn E. Holt

## Abstract

**Background:** *Klebsiella pneumoniae* is a recognised agent of multidrug-resistant (MDR) healthcare-associated infections, however individual strains vary in their virulence potential due to the presence of mobile accessory genes. In particular, gene clusters encoding the biosynthesis of siderophores aerobactin (*iuc*) and salmochelin (iro) are associated with invasive disease and are common amongst hypervirulent *K. pneumoniae* clones that cause severe community-associated infections such as liver abscess and pneumonia. Concerningly *iuc* has also been reported in MDR strains in the hospital setting, where it was associated with increased mortality, highlighting the need to understand, detect and track the mobility of these virulence loci in the *K. pneumoniae* population.

**Methods:** Here we examined the genetic diversity, distribution and mobilisation of *iuc* and *iro* loci among 2503 *K. pneumoniae* genomes using comparative genomics approaches, and developed tools for tracking them via genomic surveillance.

**Results:** *Iro* and *iuc* were detected at low prevalence (<10%). Considerable genetic diversity was observed, resolving into five *iro* and six *iuc* lineages that show distinct patterns of mobilisation and dissemination in the *K. pneumoniae* population. The major burden of *iuc* and *iro* amongst the genomes analysed was due to two linked lineages (*iuc1/iro1*, 74% and *iuc2/iro2*, 14%), each carried by a distinct non-self-transmissible IncFIB_K_ virulence plasmid type that we designate KpVP-1 and KpVP-2. These dominant types also carry hypermucoidy (*rmpA*) determinants and include all previously described virulence plasmids of *K. pneumoniae*. The other *iuc* and *iro* lineages were associated with diverse plasmids, including some carrying FII conjugative transfer regions and some imported from *E. coli*; the exceptions were *iro3*(mobilised by *ICEKp1*), and *iuc4* (fixed in the chromosome of *K. pneumoniae* subspecies *rhinoscleromatis*). *Iro/iuc* MGEs appear to be stably maintained at high frequency within known hypervirulent strains (ST23, ST86, etc), but were also detected at low prevalence in others such as MDR strain ST258.

**Conclusions:** *Iuc* and *iro* are mobilised in *K. pneumoniae* via a limited number of MGEs. This study provides a framework for identifying and tracking these important virulence loci, which will be important for genomic surveillance efforts including monitoring for the emergence of hypervirulent MDR *K. pneumoniae* strains.

## BACKGROUND

The enteric opportunistic bacterial pathogen *Klebsiella pneumoniae* imposes an increasing infection burden worldwide (1,2). These infections typically fall into one of two distinct categories; healthcare-associated (HA) infections caused by strains that are frequently multidrug-resistant (MDR), and community-associated (CA) infections arising from so-called hypervirulent strains that can cause highly invasive infections such as liver abscess but are usually drug sensitive (3,2). The antimicrobial resistance (AMR) and/or virulence determinants possessed by the associated bacteria are generally found on mobile genetic elements (MGEs) that transmit between *K. pneumoniae* cells via horizontal gene transfer (HGT) (4). These MGEs, most typically plasmids and integrative and conjugative elements (ICEs), are therefore important constituents of the accessory genome that imbue *K. pneumoniae* organisms with their distinct HA or CA clinical profiles.

It is apparent that a wide diversity of *K. pneumoniae* can cause infections in hospitalised patients (3,5,6), and that basic pathogenicity factors such as lipopolysaccharide, capsular polysaccharide, type 3 fimbriae and the siderophore enterobactin (Ent) are common to all *K. pneumoniae* and conserved in the chromosome as core genes (1,3). However enhanced virulence or ‘hypervirulence’ is associated with specific capsular serotypes (K1, K2, K5) and with MGE-encoded accessory genes that are much rarer in the *K. pneumoniae* population (3). Of particular importance are those encoding additional siderophore systems, namely yersiniabactin (Ybt) (3,7,8), aerobactin (Iuc) (9) and salmochelin (Iro)(10).

Synthesis of acquired siderophores contributes to *K. pneumoniae* virulence via multiple mechanisms. Iron assimilation via the conserved siderophore Ent is hampered by human neutrophils and epithelial cells through the secretion of lipocalin-2 (Lcn2), which binds and inhibits subsequent bacterial uptake of iron-loaded Ent (11). Ybt, Iro and Iuc are not subject to Lcn2 binding: Iro is a glycosylated derivative of Ent, while Ybt and Iuc possess an entirely distinct structure from Ent. The ability of salmochelin to counter Lcn2 binding is important for bacterial growth, and has been shown to correlate with enhanced virulence in a mouse sepsis model (12). The association between aerobactin and virulence has long been recognised, with multiple studies demonstrating its key role in increased iron acquisition, bacterial growth and/or virulence in various murine models, human ascites fluid and blood (9,13–15). Even in strains that possess all four siderophore-encoding loci, Iuc appears to play the most critical role in virulence both *in vitro* and *in vivo* (13), and serves as an important biomarker for identifying hypervirulent isolates (16).

In *K. pneumoniae*, Ybt biosynthesis is encoded by the *ybt* locus, which is typically located on a chromosomal ICE known as *ICEKp* (of which there are at least 14 distinct variants) and was recently also reported on plasmids (7,8,17). A screen of 2500 *K. pneumoniae* genomes showed *ybt* to be prevalent in one third of the population, and associated with hundreds of ICEKp acquisition events across the chromosomes of both hypervirulent and MDR lineages (8). In contrast, Iuc and Iro synthesis is encoded by loci (*iuc* and *iro*, depicted in **Fig. 1**) that are typically colocated on the so-called “virulence plasmids” of *K. pneumoniae*. The best characterised virulence plasmids are the 224 kbp plasmid pK2044 from serotype K1, sequence type (ST) 23 strain NTUH-K2044 (18); the 219 kbp plasmid pLVPK from K2, ST86 strain CG43 (19); and the 121 kbp plasmid Kp52.145pII from serotype K2, ST66 strain Kp52.145 (strain also known as 52145 or B5055; plasmid also known as pKP100) (9,20). These plasmids also carry additional virulence determinants including *rmpA* genes that upregulate capsule production, conferring a hypermucoid phenotype that is considered a hallmark of hypervirulent strains (21); other gene clusters associated with iron uptake and utilisation; and other loci encoding resistance to heavy metals such as copper (*pco-pbr*), silver (*sil*) and tellurite (*ter*) (4). In addition to the virulence plasmid-encoded *iro* and *rmpA* genes, the ST23 strain NTUH-K2044 also carries a chromosomal copy of *iro* and *rmpA* located within *ICEKp1* (7); however this is not a typical feature of ST23 (22).

**Figure 1.**
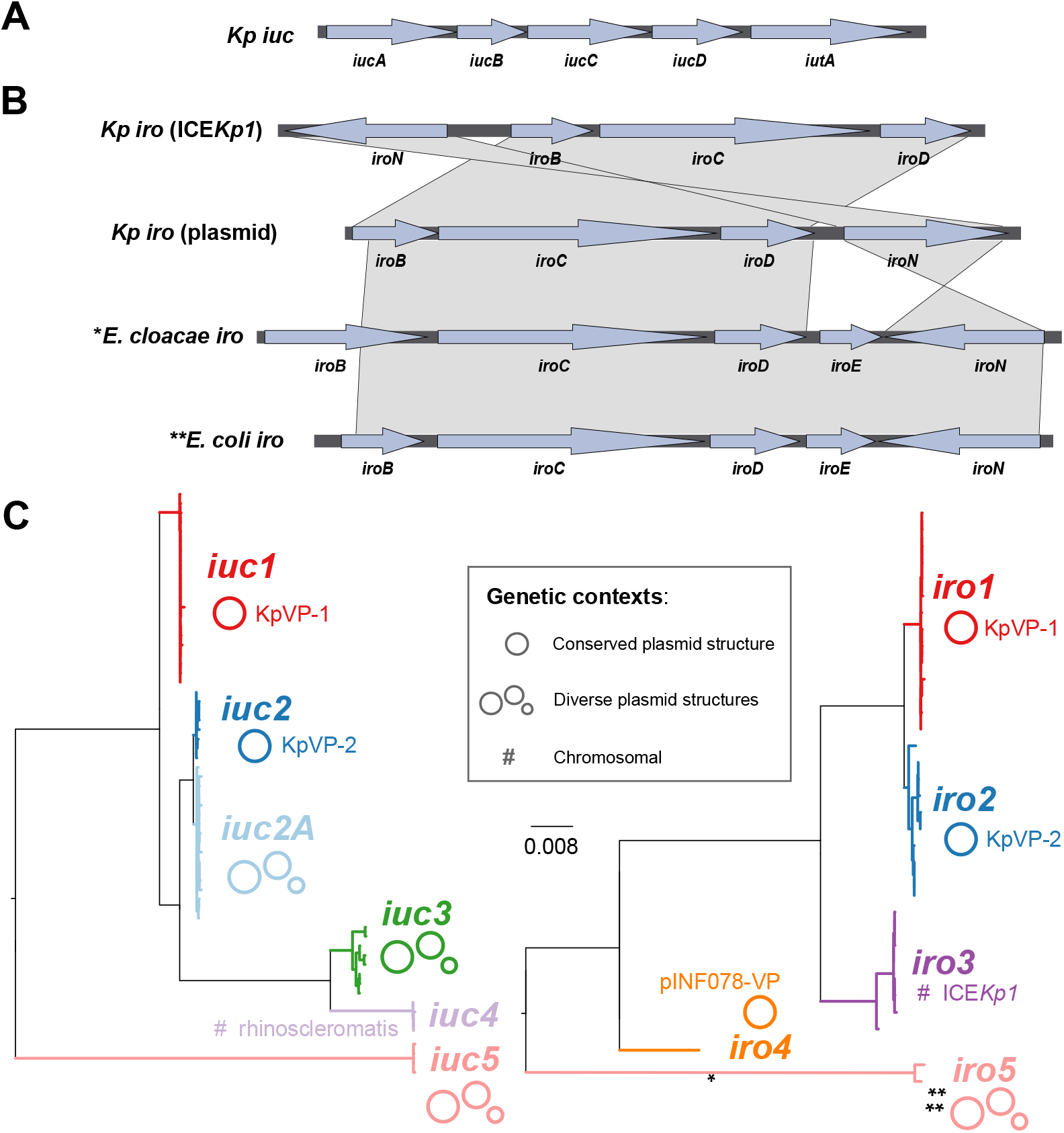
(**a**) Structure of the aerobactin locus *iuc* found in *Klebsiella pneumoniae*.(**b**) Structure of the salmochelin (*iro*) loci found in *K. pneumoniae* integrative conjugative element ICE*Kp1, K pneumoniae* plasmids, *Enterobacter cloacae* and *Escherichia coli*. *Also found in *iro4 K. pneumoniae* plasmid; **also found in *iro5 Kp* plasmids. (**c**) Maximum likelihood phylogenetic trees inferred from *iuc* and *iro* sequence types (AbSTs and SmSTs) identified in *K. pneumoniae* genomes. Phylogenetic lineages discussed in the text are labelled and their mobility indicated; nucleotide divergence within and between lineages is given in Tables S7 and S8. * and ** indicate *iro* locus structural variants that are more typical of *non-Klebsiella* species, as shown in panel (**b**).

The majority of *K. pneumoniae* lineages associated with liver abscess and other invasive community acquired infections (e.g. clonal group (CG) 23, CG86, CG380) carry virulence plasmids encoding *iro, iuc* and *rmpA* (3,9,23–25,16). However whilst virulence and AMR genes are both transmitted within the *K. pneumoniae* population via plasmids, until recently these plasmids have mainly been segregated in non-overlapping populations such that the virulence plasmids encoding *iuc* and *iro* have rarely been detected in MDR populations that cause HA infections and outbreaks (4,3,26). However, the virulence plasmid Kp52.145pII has been shown experimentally to be mobilisable (21), and there are emerging reports of MDR clones such as ST11, ST147 and ST15 acquiring virulence plasmids (27,28). The combination of hypervirulence and MDR can result in invasive infections that are very difficult to treat. This can result in dangerous hospital outbreaks; for example an aerobactin-producing carbapenemase-producing ST11 strain recently caused a fatal outbreak of ventilator-associated pneumonia in a Chinese intensive care unit, with 100% mortality (27,29). AMR plasmids are also occasionally acquired by ST23 and other hypervirulent *K. pneumoniae* clones (25,30,31).

The ease with which virulence plasmids spread in the *K. pneumoniae*population poses a significant global health threat, highlighting the importance of understanding and monitoring the movement of these loci between different strains and clones. Here we investigate the diversity of aerobactin and salmochelin synthesis loci in 2733 *K pneumoniae* complex genomes, aiming to understand the diversity and distribution of these virulence loci in the population, and to develop a framework for their inclusion in genomic surveillance efforts.

## METHODS

### Bacterial genome sequences

2733 genomes of the *K. pneumoniae* complex, including isolates collected from diverse sources and geographical locations were analysed in this study (see **Table S1**). The genomes represent a convenience sample of our own isolate collections from clinical and species diversity studies (5,8,22,32), as well as sequences that were publicly available in GenBank or via the NCTC 3000 project (https://www.sanger.ac.uk/resources/downloads/bacteria/nctc/) at the commencement of the study. The majority of these genomes were also included in our previous genome study screening for yersiniabactin and colibactin (8).

For n=1847 genomes (see **Table S1**), Illumina short reads were available and these were used to generate consistently optimised *de novo* assembly graphs using Unicycler v0.3.0b with SPAdes v3.8.1 (33,34). The remaining n=886 genomes were publicly available only in the form of draft genome assemblies. All genome assemblies were re-annotated using Prokka (35) to allow for standardised comparison. All genomes were assigned to species by comparison to a curated set of Enterobacteriaceae genomes, using mash (implemented in Kleborate, https://github.com/katholt/Kleborate); this confirmed 2503 *K. pneumoniae*, 12 *K. quasipneumoniae* subsp. *quasipneumoniae*, 59 *K. quasipneumoniae* subsp. *similipneumoniae*, 158 *K. variicola* and 1 *K. quasivariicola* (**Table S1**).

### Long read sequencing of isolates

Three isolates in our own collection (INF078, INF151, INF237) carried novel *iuc* and/or *iro* plasmids identified from short read Illumina data. We subjected these to long read sequencing using a MinION R9.4 flow cell (Oxford Nanopore Technologies (ONT)) device in order to resolve the complete sequences for the relevant plasmids. Overnight cultures of each isolate were prepared in LB broth at 37°C, and DNA extracted using Agencourt Genfind v2 (Beckman Coulter) according to a previously described protocol (doi:`10.17504/protocols.io.p5mdq46`). Sequencing libraries were prepared using a 1D Ligation library (SQK-LSK108) and Native Barcoding (EXP-NBD103) as previously described (22,36). The resulting reads were combined with their respective Illumina reads to generate a hybrid assembly using our Unicycler software v0.4.4-beta (33,36). Annotations for the assemblies were generated as described above, and the annotated sequences submitted to GenBank under accession numbers TBC (**Table S1, Table S2, Table S3**).

### Multi-locus sequence typing (MLST)

Chromosomal sequence types were determined for each genome assembly using the BIGSdb-*Kp* seven-locus MLST scheme (37) screened using Kleborate (https://github.com/katholt/Kleborate). A novel ST (ST3370) was identified and added to the BIGSdb-Kp MLST database.

To facilitate the development of MLST schemes for the aerobactin and salmochelin biosynthesis loci *iuc* and *iro*, alleles for genes belonging to each locus (i.e. *iucABCD, iutA*; and *iroBCDN;* respectively) from strains with ‘typeable’ loci (defined as those in which all genes in the locus had high quality consensus base calls when mapping with SRST2) were extracted by comparison to known alleles in the BIGSdb-Kp virulence database (http://bigsdb.pasteur.fr/klebsiella/klebsiella.html) (25), using SRST2 v0.2.0 (38) to screen Illumina read sets where available and BLAST+ v2.2.30 to screen assemblies. Incomplete, ‘non-typeable’ *iro* and *iuc* loci were excluded from the MLST scheme (marked NT in **Table S1**). Each unique combination of alleles was assigned an aerobactin sequence type (AbST) or salmochelin sequence type (SmST), defined in **Table S4 and Table S5**. The AbST and SmST schemes, profiles and corresponding alleles are also available in the BIGSdb-Kp database and in the Kleborate Github repository (see links above).

### Identification of other genes of interest, and genetic context of *iuc* and *iro* loci

Capsule (K) loci were identified in the assembled genomes using Kaptive (39). *RmpA* gene copy number was determined by BLASTn search of all genome assemblies using the *rmpA* and *rmpA2* sequences from pK2044 (GenBank accession AP006726.1) as queries. Similarly, BLASTn was used to screen the genomes assemblies for the IncFIB_K_ *repA* sequence from virulence plasmids pK2044 and Kp52.145 pII (GenBank accession FO834905.1), with FIB_K_ presence defined as >90% coverage and >80% nucleotide identity to these query sequences. FII replicons were identified using BLASTn search of the PlasmidFinder database (40).

Assemblies of all *iuc+* or *iro+* genomes were manually inspected to determine whether the loci of interest were located on the chromosome or on previously described virulence plasmids (pK2044 and Kp52.145pII). This confirmed most to be located in the chromosome (*iro3* in *ICEKp1*, or *iuc4* in the subspecies *rhinoscleromatis* lineage) or one of the known plasmids. For the remaining genomes, annotated contigs containing the *iuc* and/or *iro* loci were checked for known chromosomal or plasmid features, aided by BLASTn searching against the NCBI non-redundant nucleotide database and inspection of the assembly graphs using Bandage v0.8.0 (41).

### Phylogenetic analyses

Maximum likelihood phylogenetic trees capturing the relationships between AbSTs or SmSTs were constructed by aligning the allele nucleotide sequences corresponding to each sequence type within each scheme using MUSCLE v3.8.31 (42), then using each of the two alignments (one for AbSTs, one for SmSTs) as input for phylogenetic inference in RAxML v7.7.2 (43). For each alignment, RAxML was run fives times with the generalised time-reversible model and a Gamma distribution, and the trees with the highest likelihood were selected. Lineages were defined as monophyletic groups of AbSTs or SmSTs associated with the same MGE structure; STs within lineages shared ≥2 alleles (for SmSTs) or ≥3 alleles (for AbST), whereas no alleles were shared between lineages.

Maximum likelihood phylogenies were similarly constructed for (i) aerobactin and salmochelin locus alignments populated by sequences extracted from BLAST hits amongst representatives of the wider Enterobacteriaceae family (representatives listed in Table S6), and (ii) FIBK replicon sequence alignments constructed by mapping *iuc* positive (*iuc+*) and *iro* positive (*iro+*) genomes to a reference IncFIBK sequence (coordinates 128130 to 132007, spanning *repA* to *sopB*, of the pK2044 plasmid sequence; GenBank accession AP006726.1).

### Plasmid comparisons

Twelve representative plasmids (10 complete and 2 partial) were chosen for comparative analysis (these are available as a set in FigShare under doi:10.6084/m9.figshare.6839981; and see Table S2 for list of sources and GenBank accession numbers). Six of these representative plasmids were sourced from the NCTC 3000 project (https://www.sanger.ac.uk/resources/downloads/bacteria/nctc/). The representative plasmid sequences were compared using Mauve v2.4.0 (44), in order to identify homology blocks conserved amongst subsets of the plasmids. BLASTn comparisons of related plasmids were plotted using GenoPlotR v0.8.7 package (45) for R. All iuc+ or iro+ genomes were mapped against all 12 representative plasmids in order to calculate coverage of each plasmid in each genome. This was done using Bowtie2 v2.2.9 (46) to map Illumina reads where available, and 100 bp reads simulated from draft assemblies where raw sequence reads were not available, using the RedDog pipeline (https://github.com/katholt/RedDog). For every gene annotated within each reference plasmid, the proportion of strains within each group of genomes sharing the same *iuc/iro* lineage carrying the gene was calculated using the gene presence/absence table reported by RedDog (presence defined as ≥95% of the length of the gene being covered by at least five reads), and plotted as circular heatmaps using ggplot2 in R (using geom_tile to achieve a heatmap grid and polar_coord to circularize).

## RESULTS

### Prevalence of *iuc* and *iro* in *K. pneumoniae*

*Iuc* and *iro* were detected only in *K. pneumoniae* genomes, and not in other members of the *K. pneumoniae* species complex. Of the 2503 *K. pneumoniae* genomes screened, *iuc* was detected in 8.7% (n=217) and *iro* in 7.2% (n=181; listed in Table S1, excluding strains with a partial *iro* locus as discussed below). The presence of intact *iro* and *iuc* loci was strongly associated (odds ratio (OR) 711, 95% confidence interval (CI) 386-1458, p<1×10^-16^), co-occurring in 162 genomes (6.5% of the genomes tested). The *iro* locus appears to be susceptible to deletion: partial *iro* loci were observed in n=50 *K. pneumoniae* isolates (noted as *iro*\* in **Table S1**), mostly those that were isolated from historical collections prior to 1960 (of 39 strains isolated up to 1960 and with any *iro* genes present, 36 (92%) carried deletion variants of the locus, compared to 4/163 (2.5%) amongst isolates from 1975 onwards; OR 416, 95% CI 88-3297, p<2×10^-16^). As expected, the presence of *iuc* and *iro* were each strongly associated with presence of *rmpA*, with 157 genomes carrying all three loci (excluding partial *iro)*. A total of 238 strains (9.5%) carried *rmpA* genes: n=110 (4.4%) carried one, n=127 (5.1%) carried two, and a single strain, ST23 NTUH-K2044, carried three (as described previously (7,18), see **Table S1**).

### Genetic diversity of *iuc* and *iro* in *K. pneumoniae*

Next we explored nucleotide diversity of the genes comprising the *iro* and *iuc*loci in *K pneumoniae*. The five genes comprising the *iuc* locus (**Fig. 1a**) and four genes of the *K. pneumoniae* form of the *iro* locus (**Fig. 1b**) were screened for sequence variation, and each unique locus sequence variant was assigned an allele number. Of the n=209 strains carrying a typeable *iuc* locus, 62 unique *iuc* allele combinations were observed and assigned a unique aerobactin sequence type or AbST (see **Table S4** for AbST definitions, and **Table S1** for AbSTs assigned to each genome). The *iutA* alleles present in the *iuc* locus showed >28% nucleotide divergence from the core chromosomal homolog of *iutA* encoding a TonB-dependent siderophore receptor, which was detected in 96.4% of all genomes; the alleles of this core chromosomal gene are not included in the aerobactin MLST scheme. Typeable *iro* loci were identified in n=164 strains, comprising 35 unique salmochelin sequence types or SmSTs (defined in **Table S5**, see **Table S1** for SmSTs assigned to each genome). Maximum likelihood phylogenetic analyses of the AbST and SmST sequences, and their translated amino acid sequences, revealed five highly distinct *iuc* lineages and five *iro* lineages (labelled *iro1, iro2* etc; see **Fig. 1c, Fig. S1**). Nucleotide divergence between lineages was 1-11% (20-1000 substitutions), and no alleles were shared between lineages (**Table S7, S8**). Nucleotide divergence within lineages was low, with mean divergence of 0.001-0.40%(*iro*) and 0.013-0.50%(*iuc*) (**Table S7, S8**), and at least two (*iro*) or three (*iuc*) shared alleles between members of the same lineage. Of note, the *iro4, iro5* and *iuc5* loci were quite distant from other lineages (each showing >5.5% nucleotide divergence from all other lineages, vs <4.6% divergence amongst the other lineages; **Fig. 1, Table S7, S8**). Comparison to *iuc* and *iro* genes present in other Enterobacteriaceae (see **Fig. S2, Table S9**), and the presence of the additional *iroE* gene (**Fig. 1b**), suggests that these more distant lineages derive from outside *Klebsiella*, most likely *Enterobacter* (*iro4*) and *E. coli* (*iro5, iuc5*). Note that genotyping of *rmpA* was not performed since most *rmpA*-positive genomes carry two copies of the gene, which complicates allele typing from short read data, however *rmpA* copy number per genome is reported in **Table S1**.

### Mobile genetic elements associated with *iuc* and *iro* loci

Inspection of the genetic context surrounding the *iuc* and *iro* sequences revealed that the various *iuc* and *iro* lineages were associated with distinct MGEs, with the exception of *iuc4* which was restricted to the chromosome of *K. pneumoniae*subspecies *rhinoscleromatis* (ST67) (**Fig. 1c, Table 1**). Most common were *iuc1* and *iro1;* these were both associated with pK2044-like plasmids (hereafter called KpVP1-1, see below) and the presence of two *rmpA* genes, and accounted for 74% of all *iuc+iro+* genomes. These were followed by *iuc2* and *iro2*, which were associated with Kp52.145 pII-like plasmids (hereafter called KpVP-2, see below), the presence of one *rmpA* gene, and accounted for 14% of all *iuc+iro+* genomes. A sister clade of *iuc2*,which we named *iuc2a*, was associated with diverse plasmids that shared some homology with Kp52.145 pII (36-70% coverage, 99% nucleotide identity). Most *iuc2a+* isolates carried a single *rmpA* gene (n=38, 88.4%) and all lacked an intact *iro* locus (n=26, 60.5% had a partial *iro* locus). Lineage *iuc3* was related to the *iuc4* lineage encoded on the *rhinoscleromatis* chromosome, but was present on novel plasmids. *Iro3* was located within the chromosomally integrated *ICEKp1*, along with *rmpA*. Four genomes carried *iuc5* (two of these also carried *iro5;* all lacked *rmpA)*. The *iuc5* sequences were distantly related to *iuc1* and *iuc2* (>8.9% nucleotide divergence), but were identical to sequences found in *E. coli* and located on contigs that matched closely to *E. coli* AMR plasmids (e.g. strain PCN033 plasmid p3PCN033, accession CP006635.1 (47), which showed >99% nucleotide identity to the best assembled of *iuc5+ K. pneumoniae* contigs). *Iro4* was identified in a single genome (which lacked *rmpA*), and was >6.1% divergent from *iro1* and *iro2*sequences. Its closest known relatives are *iro* sequences present in the chromosomes of *Enterobacter cloacae* and *Enterobacter hormaechei* (strains AR_0065, accession CP020053.1 and 34977, accession CP010376.2 respectively; 95% identity). Lineages *iro4* and *iro5* follow the gene configuration typical of non-K. *pneumoniae* Enterobacteriaceae *iro* loci, from which the *K. pneumoniae iro1, iro2* and *iro3* differ by lack of *iroE* and inversion of *iroN* (see Fig. 1b).

**Table 1.**
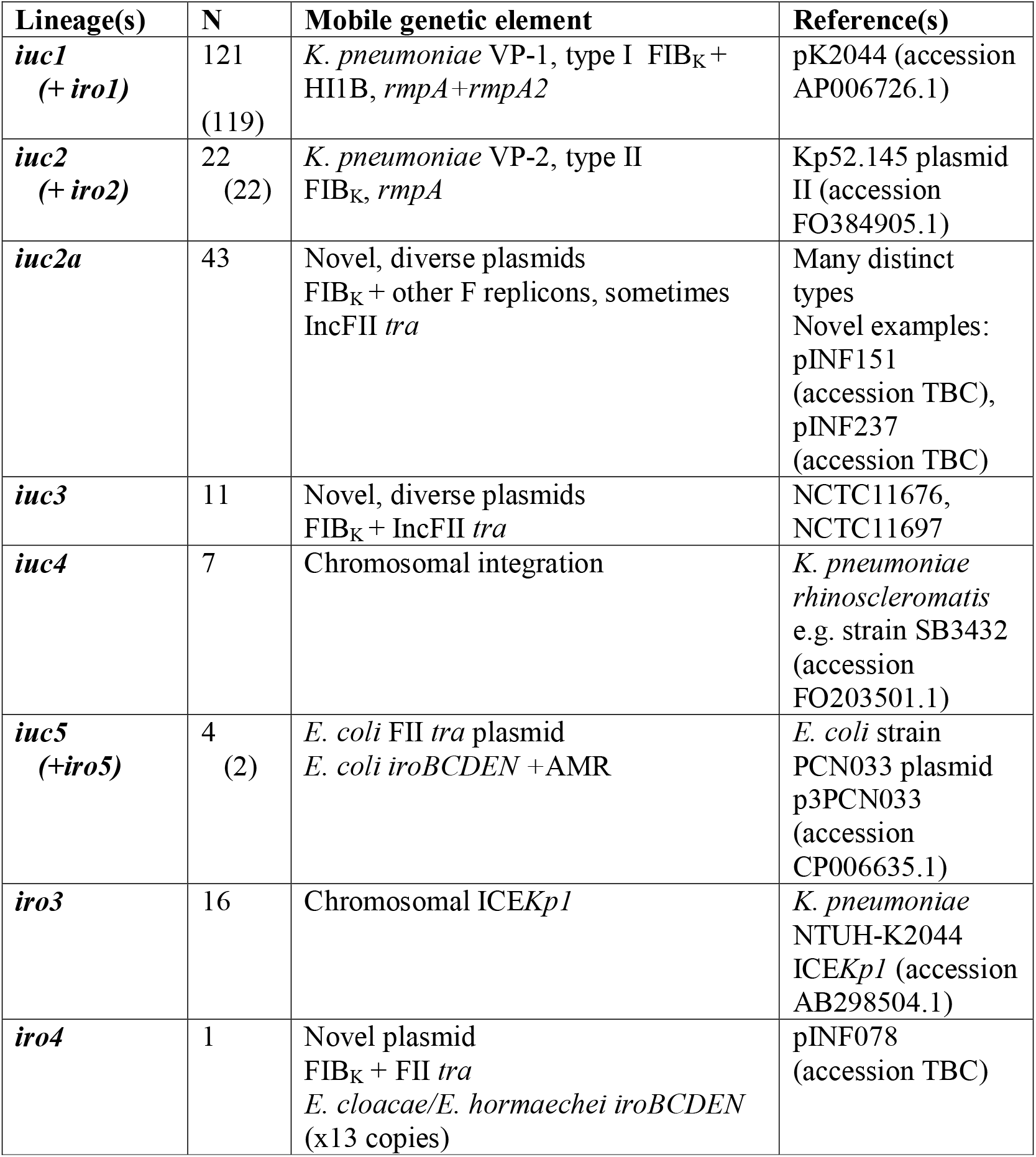
Summary of *iuc* and/or *iro* plasmid lineages

To examine the gene content and replicon differences between the various *K. pneumoniae* plasmids associated with *iuc* and/or *iro*, 12 representative plasmids associated with the various lineages were selected for comparison (Fig. 2, Table S2). These include six complete *K. pneumoniae* plasmid sequences identified from finished genomes: *iuc1/iro1* (n=1), *iuc2/iro2* (n=1), *iuc2a* (n=3), *iuc3* (n=1); three novel complete *K. pneumoniae* plasmid sequences that we generated for this study, carrying *iuc2a* (n=2) and *iro4* (n=1); and 2 large contigs that we identified from public *K. pneumoniae* genome data representing partial sequences for additional plasmids carrying *iuc2a* (n=1) and *iuc3* (n=1) (**Fig. 2**). The *K. pneumoniae* genomes in which *iuc5/iro5* were identified were available only as draft assemblies deposited in public databases, and the associated plasmid sequences were fragmented in these assemblies, hence we used *E. coli* strain PCN033 plasmid p3PCN033 (47) as the representative for *iuc5*/*iro5*. The representative plasmid sequences differed substantially in their structure and gene content (**Fig. 2**), and were differentially distributed amongst the *K. pneumoniae* population (**Fig. 3**). In order to explore structural conservation of plasmids amongst isolates with each *iro* or *iuc* lineage, we mapped the sequence data from all isolates carrying either of these loci against the 12 representative plasmid sequences (**Fig. 4**). This revealed that plasmid structures were largely conserved amongst isolates sharing the same *iuc* or *iro* lineages, although plasmids associated with *iuc2a* and *iuc3* showed more diversity than others (**Fig. 4** and see below).

**Figure 2.**
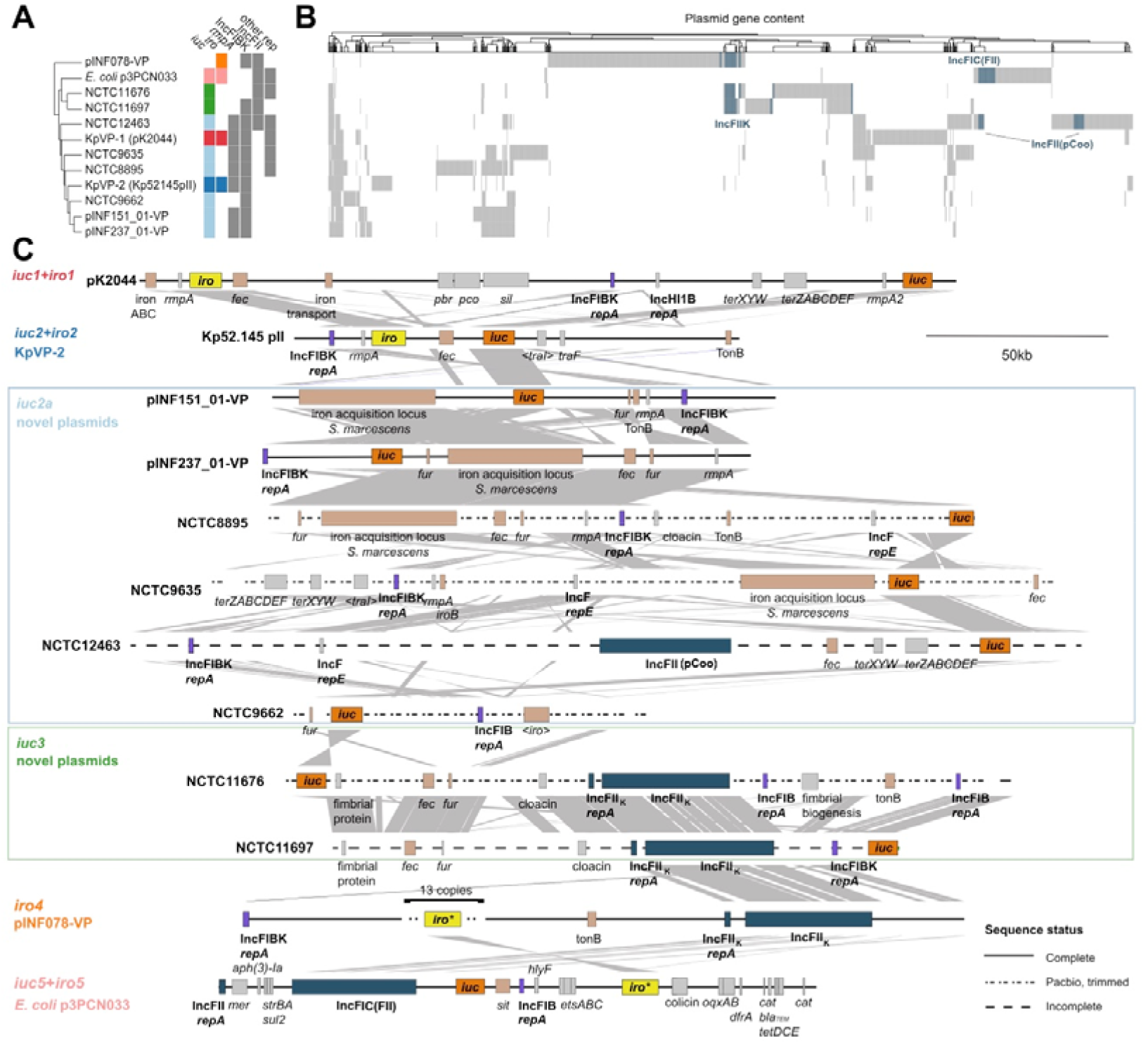
Plasmid variants associated with different *iro* and/or *iuc* lineages identified amongst *K. pneumoniae*. (**a**) Clustering of the 12 reference plasmids based on gene content, annotated with presence of *iuc* and *iro* lineages (coloured as in panel **b**, and **Figure 1c**), *rmpA*, IncFIB_K_, IncFIB, IncFII and/or other plasmid replicon types. (**b**) Gene content matrix for reference plasmids; columns correspond to protein-coding sequences that are >10% divergent from one another. IncFII conjugal transfer region genes are coloured blue, to highlight the divergent forms of this region and labelled with the closest IncFII type as detected by PlasmidFinder. (**c**) Genetic maps for the reference plasmids. The positions of key loci involved in core plasmid functions (**bold**), virulence (*iro* highlighted in yellow, *iuc* in dark orange and other loci involved in iron acquisition/transport in light orange) and antimicrobial resistance are indicated. Grey shading indicates homology blocks sharing >60% nucleotide identity.

**Figure 3.**
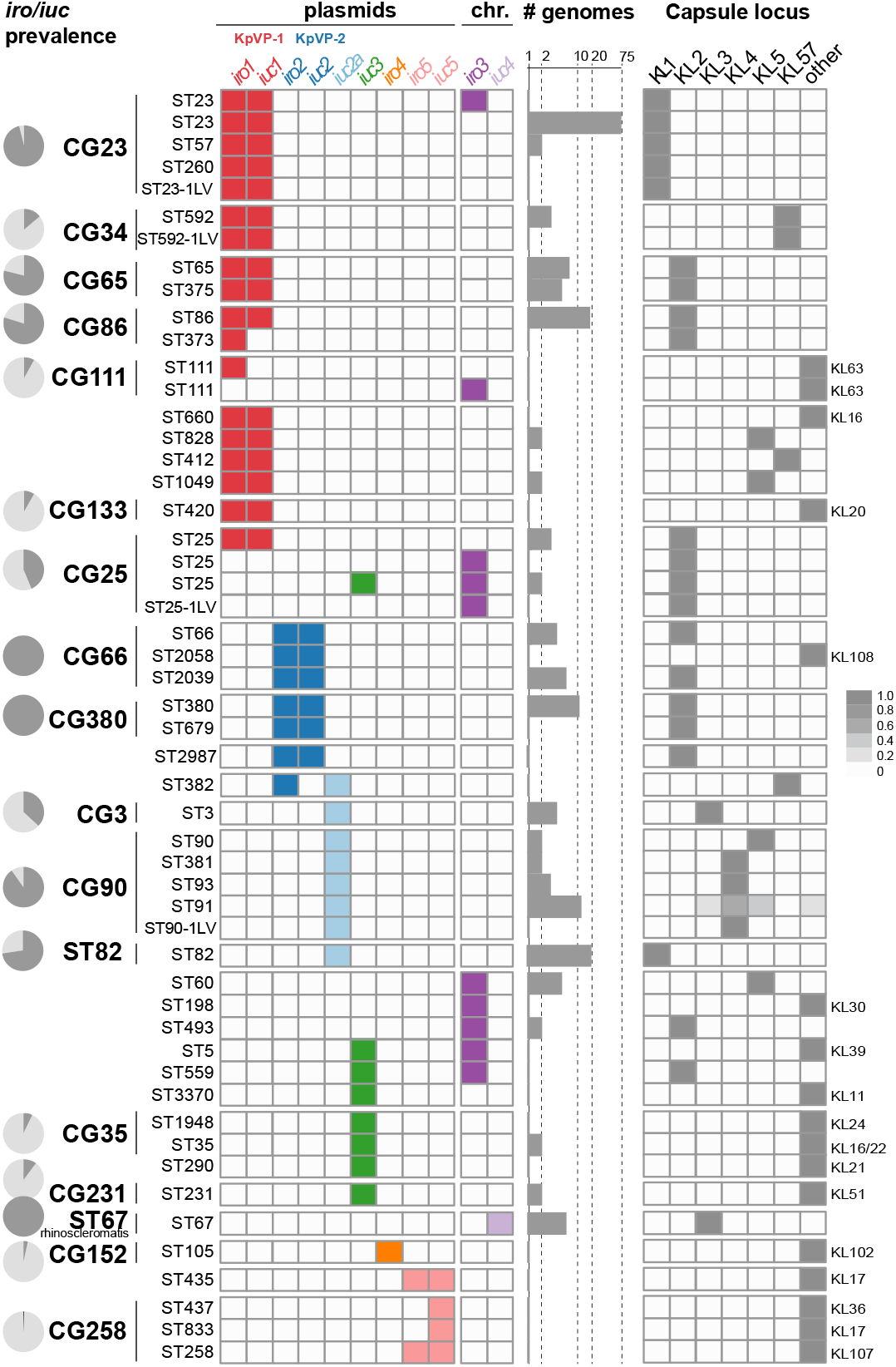
Distribution of plasmid and chromosomal variants of *iro* and *iuc* and capsule locus (KL) types amongst *K. pneumoniae* clones. Rows indicate sequence types (STs, as labelled) that contain ≥1 genome in which *iro* and/or *iuc* was detected; vertical lines indicate STs belonging to the same clonal group (CG) as labelled. Pie charts indicate prevalence of *iro* and/or *iuc* within common *K. pneumoniae* lineages. The detection of individual *iro* and *iuc* lineages within each *K. pneumoniae* ST is indicated in the grid, coloured as per **Figure 1**. Bar plots indicate sample size (number of genomes per ST; note log10 scale). Heatmap on the right indicates prevalence of capsule (K) locus types in each *K. pneumoniae* ST, coloured as per inset legend. Individual columns are included for K types that are common amongst virulent clones; where other K types were detected these are represented in the ‘other’ column, and the relevant K type for that ST is labelled to the right.

**Figure 4.**
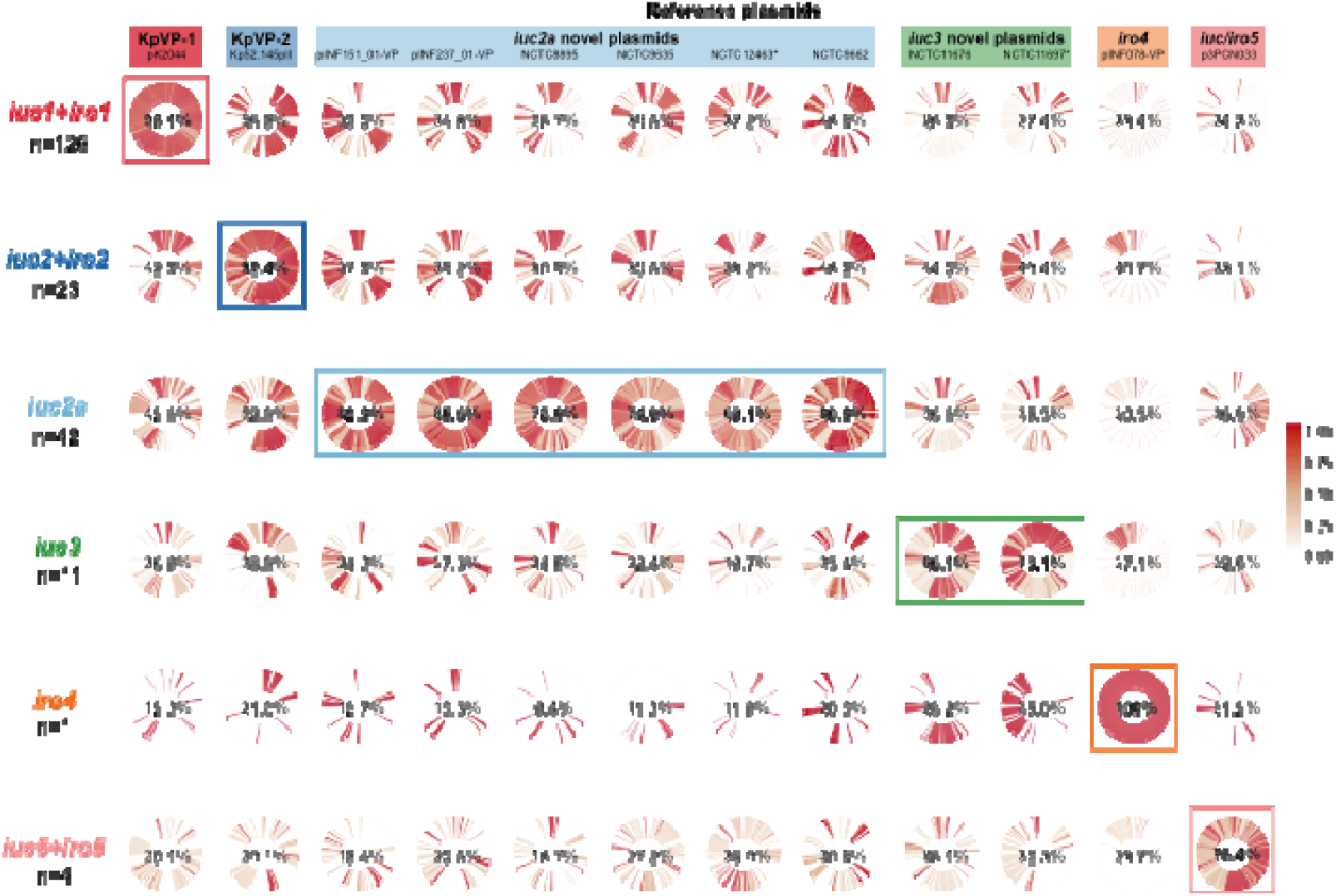
Conservation of reference plasmid genes amongst strains with plasmid-associated *iuc/iro* lineages. Cells show circularised heatmaps indicating the frequency of each gene in a given reference plasmid (column); amongst strains that contain a given *iro* and/or *iuc* lineage (row). Around each circle, genes are ordered by their order in the corresponding reference plasmid. Percentages in the middle of each cell indicates the mean coverage of the reference plasmid sequence (column), amongst strains belonging to each *iro/iuc* lineage (row); bold labels and boxes highlight groups of strains carrying the same *iuc/iro* lineage as the reference plasmid. * indicates the two plasmids represented by incomplete plasmid sequences.

All representative *iuc* or *iro* plasmids harboured an IncFIBK (n=9) or IncFIB (n=3) replicon, including the *repA* replication gene, *oriT* origin of transfer, and *sopAB* partitioning genes (presence of these replicons in each plasmid is indicated purple in **Fig. 2c**, and listed in **Table S2**). The IncFIBK replicon was present in n=202/208 (97%) of isolates with plasmid-encoded *iuc* or *iro*, including 100% of *iuc1/iro1, iuc2/iro2, iuc2a* and *iro4* isolates; and 82% of *iuc3* isolates. Each of these *iuc/iro* lineages was associated with a unique sequence variant of the IncFIB_K_ replicon (see tree in **Fig. 5** and nucleotide identity with the IncFIBK *rep* sequences from KpVP-1 and KpVP-2 listed in **Table S1**), supporting segregation of the *iuc* and *iro* loci with distinct FIBK plasmid backbones. However the IncFIBK replicon was also widely detected amongst strains that do not carry *iro* and *iuc* (77% of all *K. pneumoniae* genomes and 69% amongst other species in the complex; see **Table S1**), including MDR *K. pneumoniae* lineages such as CG258, and is known to be associated with AMR plasmids (48,49). IncFIB replicons, which are common amongst *E. coli* and display >39% nucleotide divergence from the FIB_K_ replicon, were found in all *K. pneumoniae* strains carrying the *E. coli* variant *iuc5* (100%) and also detected in two strains carrying *iuc3* plasmids (18%; marked in **Fig. 2**), suggesting transfer of these *iuc* variants into *K. pneumoniae* via such plasmids.

**Figure 5.**
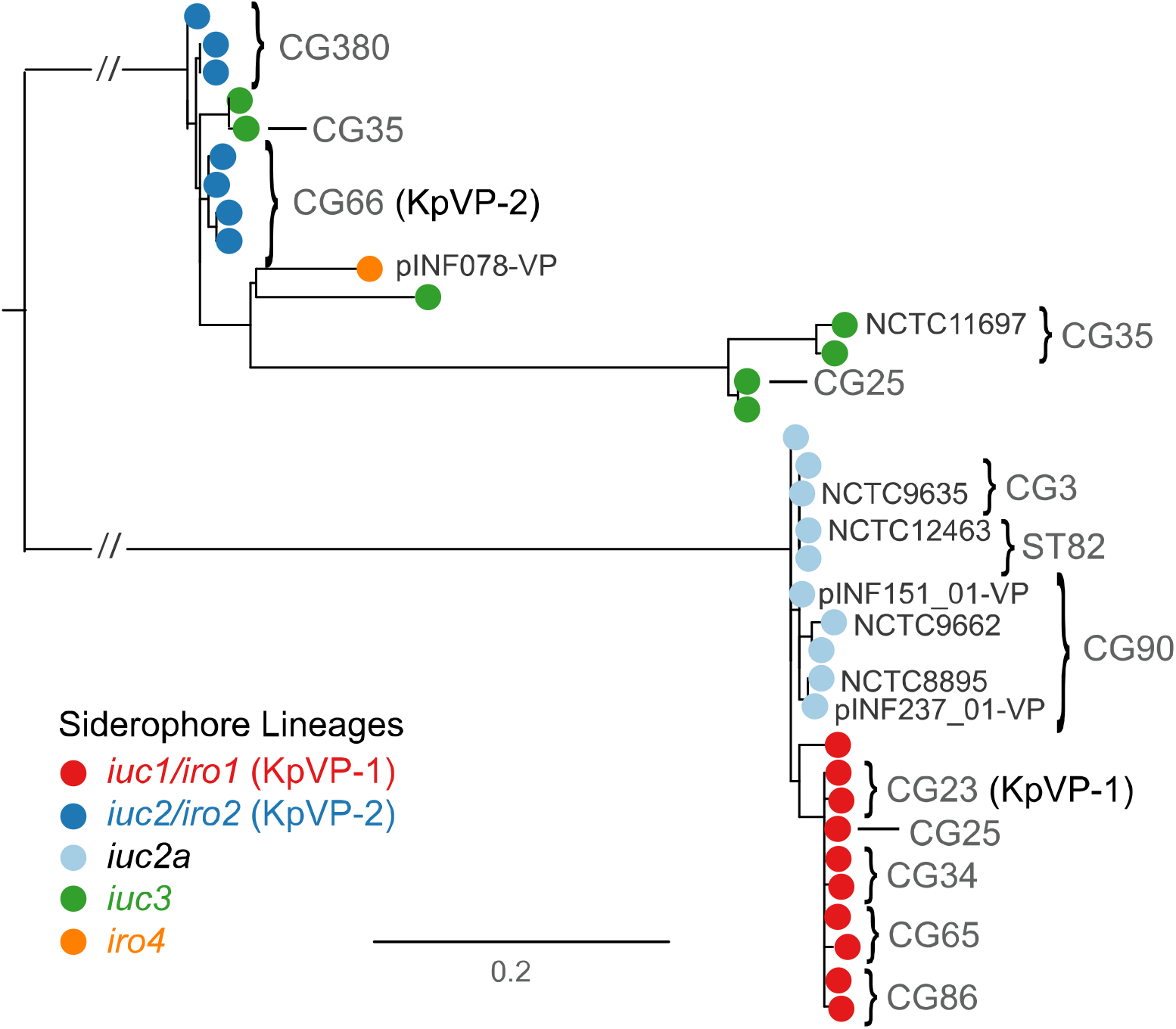
Maximum likelihood phylogeny of representative IncFIB_K_ replicon sequences from strains with *iuc/iro* plasmids. Each tip represents a unique IncFIB_K_ replicon sequence (spanning *repA, oriT, sopAB*), coloured according to the *iro/iuc* lineage carried by the corresponding strains as per inset legend. FIBK sequences found in the representative plasmid sequences (shown in **Fig. 2** and listed in **Table S2**) are labelled; tips/subclades are also annotated to indicate those found in common clonal groups (CG; see **Fig. 3**).

### *Iuc1 iro* lineages 1 and 2 are associated with two dominant *K. pneumoniae* virulence plasmids, KpVP-1 and KpVP-2

*Iuc/iro* lineages 1 and 2 accounted for 64% of *K. pneumoniae* isolates carrying any aerobactin or salmochelin synthesis loci, and 88% of isolates carrying both. Whilst it was not possible to resolve the complete sequences for all plasmids associated with these lineages, read mapping to pK2044 and Kp52.145 pII reference sequences strongly supported the presence of pK2044-like plasmids in *iro1+iuc1+*genomes (mean plasmid coverage of 95.1%, range 28.8-100%; see **Fig. 4**), and Kp52.145 pII-like plasmids in *iro2+iuc2+* genomes (mean plasmid coverage of 92.4%, range 87.2-100%; see **Fig. 4**). There were limited homologous regions shared between the two plasmids (**Fig. 2**), including the *iro, iuc, rmpA* and *fec* loci, and the IncFIBK replicon (Table S10). These shared regions were largely conserved across all isolates carrying *iuc/iro* lineages 1 or 2; the remaining regions unique to either pK2044 or Kp52.145 pII were largely conserved amongst the isolates that carried lineage 1 or 2 loci, respectively (**Fig. 4**). Notably, the loci encoding heavy metal resistances against copper (*pbr-pco*), silver (*sil*) and tellurite (*terXYW* and *terZABCDEF*) were highly conserved amongst lineage 1 strains but not present in any of the lineage 2 strains (**Table S10**). As noted above, *iuc/iro* lineages 1 and 2 were also each associated with a distinct variant of the IncFIBK replicon sequence (**Fig. 5**). Hence we define pK2044-like plasmids carrying *iuc1* and *iro1* loci as *K. pneumoniae* virulence plasmid type 1 (KpVP-1), with reference plasmid pK2044; and Kp52.145 pII-like plasmids carrying *iuc2* and *iro2* loci as *K. pneumoniae* virulence plasmid type 2 (KpVP-2). Both plasmid types typically carry at least one copy of *rmpA;* neither one carries genes associated with conjugation, hence we assume they are not self-transmissible.

KpVP-1 and KpVP-2 showed distinct distributions within the *K. pneumoniae* population. KpVP-1 was present in 5.0% of all isolates and accounted for 74% of *iuc+iro+* isolates. The KpVP-1 reference plasmid pK2044 originated from an ST23 isolate (CG23), and KpVP-1 was strongly associated with this and two other well-known hypervirulent clones CG65 and CG86, in which it was present at high prevalence (ranging from 79.0-96.4%, see **Fig. 3**). KpVP-1 was also detected at low frequencies in other clones, including CG34, CG111, CG113 and CG25, suggesting it is mobile within the *K. pneumoniae* population (**Fig. 3**). KpVP-2 was present in 0.96% of all isolates and accounted for 14% of *iuc+iro+* isolates. The KpVP-2 reference plasmid Kp52.145 pII originated from an ST66 isolate, and KpVP-2 was present in all isolates of the associated clonal group CG66 (n=11) and also all isolates of CG380 (n=12) (**Fig. 3**).

### An *iuc* lineage 2 variant (*iuc2a*) is associated with diverse plasmids with a KpVP-1-like IncFIBK replicon

*Iuc2a* was identified in 43 isolates largely belonging to three clonal groups (ST3, n=4; CG90, n=19; ST82, n=19; ST382, n=1; see **Fig. 3**), with the majority (n=38, 88.4%) from the historical NCTC or Murray collections and isolated between 1932 and 1960 (**Table S1**). Provenance information was available for only twelve of the *iuc2a+* isolates (1 ST3, 9 CG90, 2 ST82); all of which originated from the human respiratory tract (3 nose, 1 throat, 7 sputum, and 2 NCTC isolates recorded only as respiratory tract). We used long-read sequencing to resolve plasmids in two novel *iuc2a+* isolates from our own collection, INF151and INF237, which were both CG90 Australian hospital sputum isolates (summarised in **Table S3**). This yielded IncFIB_K_ plasmids in each genome, of size 138.1 kbp and 133.7 kbp, respectively (accessions: pINF151_01-VP, TBC; pINF237_01-VP, TBC). Both plasmids carried *iuc2a* and one *rmpA* gene, but they differed slightly from one another in structure and gene content, and differed substantially from the three complete *iuc2a+* plasmid sequences available from NCTC isolates (ST3 and CG90; see **Fig. 2, Fig. 4**). Only one of these plasmids (from NCTC 12463; incomplete) carried a conjugative transfer region (IncFII), hence we predict most are not self-transmissible. Mapping of *iuc2a+* isolates to each of the five representative *iuc2a+* plasmid sequences indicated a degree of conservation between plasmids in strains belonging to the same *K. pneumoniae* clone, but none particularly well conserved across all *iuc2a+* strains (**Fig. 4, Fig. S3**). However all *iuc2a+* isolates formed a tight monophyletic cluster in the IncFIBK replicon tree (**Fig. 5**), consistent with recent shared plasmid ancestry followed by frequent structural and gene content changes. Notably the iuc2a-associated IncFIB_K_ replicon sequences were closely related to those of KpVP-1 and distant from those of KpVP-2, hence we hypothesise that *iuc2a* plasmids share an ancestor that was a mosaic including *iuc2* -related sequences from KpVP-2 and IncFIBK replicon sequences from KpVP-1.

### *Iuc* lineage 3 is mobilised by diverse plasmids carrying the FII_K_ conjugative transfer region

Lineage *iuc3* was detected in 11 isolates from diverse sources and chromosomal STs (**Fig. 3**), and was associated with three related variants of the IncFIBK replicon (**Fig. 5**). We identified one complete and one near-complete *iuc3* plasmid sequences: a complete 189.8 kb plasmid from NCTC 11676 (isolated 1979, ST290), and a 155.4 kb contig from NCTC 11697 (isolated 1984, ST3370) (**Fig. 2**). The plasmids share around half of their gene content (96 kbp), including the IncFII_K_ *tra-trb* conjugative transfer machinery, a fimbrial protein and the *fec* iron acquisition system in addition to *iuc3* (**Fig. 2, Fig. 4, Table S2**). Mapping to these sequences showed all *iuc3+* isolates carried related plasmids with an FIIK transfer region (**Fig. 4, Table S10**).

### Complete sequence of an *ro4* plasmid

Lineage *iro4* was identified in a single hospital UTI isolate INF078 (ST105) from Australia, whose genome sequence we completed using long reads (replicons summarised in **Table S3**). Hybrid assembly using short and long reads resolved a 399,913 kbp plasmid, pINF078-VP (accession TBC) which carried multiple copies of *iro4*, the IncFIB_K_ replicon (similar to the KpVP-2 variant, see **Fig. 5**) and the IncFII_K_ replicon and transfer region (Fig. 2). As noted above, the *iro4* locus is more closely related to *Enterobacter iro* than to other *K. pneumoniae iro* in terms of both structure (including the *iroE* gene; see **Fig. 1b, Fig. S4**) and sequence (Fig. S2), suggesting it has been transferred from *Enterobacter* into a *K. pneumoniae* IncFIBK/FIIK plasmid backbone (note the IncFIBK replicon sequence of pINF078-VP was similar to those of KpVP-2; see **Fig. 5**). pINF078-VP harboured multiple tandem copies of a 17,129 bp region containing *iroBCDEN* and 12 other genes of unknown function (**Fig. S4**). Long read sequences (up to 70 kbp) spanning the non-repeat and repeat region of pINF078-VP confirmed at least n=3 copies of the 17 kbp repeated sequence, whose mean read depth in the Illumina sequence data was 13.3 times that of the rest of the plasmid sequence, suggesting approximately 13 tandem copies.

### *Iuc/iro* lineage 5 loci are associated with plasmids originating from *E. coli*

Four *K. pneumoniae* isolates carried the *E. coli* variant *iuc5;* two of these also carried the *E. coli* variant *iro5* (see species trees in Fig. S2). Three *iuc5+* isolates (including one with *iro5*) belonged to the globally-disseminated, carbapenemase-producing *K. pneumoniae* CG258 (ST258, KPC+; ST437, KPC+; ST833, KPC-) and carried several AMR genes. Unfortunately all four *iuc5+* genomes were sourced from public databases and were available in draft form only, and the complete plasmid sequences could not be resolved. However the *iuc5+* contig sequences from *K. pneumoniae* share close homology (99% identity, >65% coverage) with *iuc5+iro5+* FII conjugative plasmids from E. *coli* that also carry AMR genes (e.g. p3PCN033, CP006635.1; D3 plasmid A, CP010141.1).

## DISCUSSION

This study reveals significant genetic diversity underlying the biosynthesis of aerobactin and salmochelin in *K. pneumoniae*, but shows the distribution of *iuc* and *iro* locus variants is highly structured within the population. Our data indicate that most of the burden of these hypervirulence-associated siderophores in the *K. pneumoniae* population is associated with two dominant virulence plasmids, which we define here as KpVP-1 and KpVP-2, that differ in terms of gene content (**Fig. 2**) and are each associated with co-segregating sequences of the FIB_K_ replicon, *iuc* and *iro* loci (**Fig. 1, Fig. 5**). These dominant virulence plasmid types are each represented by one of the previously characterised *K. pneumoniae* virulence plasmids (18,20), pK2044 (KpVP-1, encoding *iro1* and *iuc1*) and Kp152.145pII (KpVP-2, encoding *iro2* and *iuc2);* both also carry hypermucoidy determinants, and together they account for 74% and 14% of the *iuc+iro+ K. pneumoniae* genomes analysed. Importantly, our data indicate that each of these common virulence plasmid variants is maintained at high prevalence in a small number of known hypervirulent clones: KpVP-1 in CG23 (96%, including pK2044 (18)), CG86 (80%, including pLVPK (19)) and CG65 (79%); KpVP-2 in CG66 (100%, including Kp152.145pII) and CG380 (100%) (**Fig. 3**). This suggests that both plasmid types can persist for long periods within a host bacterial lineage as it undergoes clonal expansion; indeed our recent study of the evolutionary history of CG23 indicates that KpVP-1 has been maintained in this clonally expanding lineage for at least a century (22). Notably we also detected KpVP-1 at low prevalence in numerous other *K. pneumoniae* lineages and KpVP-2 at low prevalence in one other lineage, suggesting the possibility of wider dissemination of both plasmid types by occasional transfer to new lineages (**Fig. 3**). Given the stability of the plasmids observed in several clonal groups, we speculate that some of these transfer events will result in the emergence of novel hypervirulent strains that can stably maintain the plasmid into the future. In contrast, the non-plasmid form of *iro* (*iro3*, occasionally integrated into the chromosomes of *K. pneumoniae* via ICE*Kp1* was found at low prevalence (<0.5%) and included just one of the 79 ST23 isolates analysed (NTUH-K2044, in which ICEKp1 was first described), 1/1 ST5, 1/21 ST111 (13%), 1/2 ST198, 2/15 CG25, 2/2 ST493 and 5/5 ST60. Hence while ICE*Kp1* is somewhat dispersed in the *K. pneumoniae* population, it shows little evidence of stability within lineages, consistent with our previous observations regarding ICE*Kp* in general (8).

We also detected several novel iuc+ or iro+ plasmid types, the most common being the group of *iuc2a* plasmids (21% of all iuc+ strains) that were detected in respiratory isolates from CG3, CG82 and CG90 and mostly originated from historical collections (50). Interestingly these combine an *iuc* sequence closely related to that of KpVP-2 (**Fig. 1**) with a FIB_K_ replicon sequence very close to that of KpVP-1 (**Fig. 5**), and showed substantial mosaicism and gene content variation (**Fig. 2, Fig. 4**). The *iuc3* lineage was also quite common (5.3% of all *iuc+* strains) and associated with a variety of diverse plasmids, most of which carried the FII conjugative transfer region and thus are likely self-transmissible (**Fig. 2, Fig. 4**). It is notable that *iuc2a* and *iuc3* plasmids were not only quite rare in the population, but also showed less evidence of stable maintenance within *K. pneumoniae* lineages (**Fig. 3**) and lower stability of gene content (**Fig. 2**) than the dominant KpVP-1 and KpVP-2 plasmids (**Fig. 4**). The position of *iuc2a* and *iuc3* in the *iuc* trees (**Fig. 1, Fig. S2**) suggests that both derive from other *K. pneumoniae* loci, hence we speculate it is the properties of the plasmids mobilising these loci, and not the siderophore biosynthesis loci themselves, that makes these variants less widespread in the *K. pneumoniae* population. This variation in gene content may be a consequence of self-transmissibility, exposing the plasmids to a wider gene pool of host bacteria and providing opportunities for gene content diversification, which could potentially include AMR genes. Notably the *iuc3* plasmids carry an arsenal of additional virulence loci involved in iron metabolism and resistance to heavy metals, reminiscent of KpVP-1 (**Fig. 2**).

The other novel plasmids appear to derive from outside *K. pneumoniae* (**Fig. 1, Fig. S2**). Most concerning are the four *E*. coli-derived plasmids we detected carrying *iuc5* (and occasionally *iro5*) in USA and Brazil, three of which were found in the MDR hospital outbreak-associated clone CG258. Whether these aerobactin plasmids harbour AMR genes as they do in *E. coli* is not currently resolvable; however it seems that conjugative *E. coli* plasmids such as D3 plasmid A do have the potential to deliver hypervirulence and multidrug resistance to *K. pneumoniae* strains in a single step. A recent study of *K. pneumoniae* submitted to Public Health England used PCR to screen for isolates carrying both carbapenemase genes and *rmpA*, as a marker of the virulence plasmid, and identified a plasmid harbouring *iuc, rmpA, rmpA2* and the AMR genes *sul1, sul2, armA, dfrA5, mph(A)* and *aph(3’)-VIb* (28). To our knowledge this is the first report of a complete sequence of a *K. pneumoniae* plasmid harbouring both AMR and virulence genes. The isolate (ST147) was not included in our original screen, however subsequent analysis using *Kleborate* plus manual inspection of the plasmid sequence reveals it carries *iuc1* (AbST63, a novel single locus variant of AbST1 which is typical of hypervirulent clones CG23, CG65 and CG86) and appears to be a mosaic carrying sequences from KpVP-1 (40% coverage), an IncFII conjugative transfer region and transposons carrying AMR genes.

The presence of aerobactin synthesis loci in the *iuc5+ K. pneumoniae* isolates we identified here was not reported in the original studies (51,52), and thus it is not known whether they actually produce aerobactin or show enhanced virulence. This highlights the need to raise awareness of the *iuc* and *iro* loci as potentially clinically relevant hypervirulence factors, and to screen for them in isolates and genome data. The latter we aim to facilitate via the genotyping schemes established here, which can be used to easily screen new genome assemblies using Kleborate (https://github.com/katholt/Kleborate/) or BIGSdb-Kp (http://bigsdb.pasteur.fr/klebsiella/klebsiella.html), or new short read data sets using SRST2 (https://github.com/katholt/srst2). PCR primers suitable for screening for *iro* and *iuc* can be found in Lee *et al* (53). Notably many studies rely on the hypermucoidy phenotype to identify hypervirulent strains, however this is dependent on growth conditions (54), and recent studies indicate that aerobactin synthesis is a more important virulence determinant (16,13,14). Our data suggest that hypermucoidy screening would typically pick up most of the common aerobactin plasmids KpVP-1, KpVP-2 and *iuc2a+* plasmids; but not those carrying *iuc3*, or the *iuc5* plasmids from *E. coli*. Additionally, it is important not to conflate the presence of the core chromosomal receptor gene *iutA* with the ability to synthesise aerobactin, which is encoded in the *iuc* locus (6). False positive detection of the aerobactin locus version of *iutA* can be avoided by using an identity threshold of <20% divergence. Tellurite resistance has also been suggested as a phenotypic screen to identify hypervirulent isolates of CG23, CG65 and CG86(55); our data confirms this is a good marker for KpVP-1 (92.6% carry *ter*), but not for other aerobactin plasmid types (Table S10).

## CONCLUSIONS

Our results illuminate that distinct virulence plasmid variants are associated with the various hypervirulent *K. pneumoniae* lineages, but also highlight that these alongside other plasmids and MGEs can shuttle aerobactin and salmochelin synthesis loci to other lineages, threatening the emergence of novel hypervirulent strains. Indeed, reports of MDR clones acquiring *iuc* plasmids appear to be increasing in incidence, particularly in China (27,29,56-58), and have been associated with increased morbidity and mortality. The AbST and SmST typing schemes developed in this study provide an important resource to identify and monitor the movement of *iro* and *iuc* loci and associated MGEs in *K. pneumoniae* genomes; which will be important to detect and contain these emerging threats. Genotyping with our tools reveals the *iuc* plasmid identified in the recently reported fatal hospital outbreak of carbapenemase-producing ST11 in Beijing is a variant of KpVP-1 that carries *iuc1* (AbST1) and a single copy of *rmpA* but lacks the *iro* locus (27). In this strain the aerobactin plasmid does not carry any AMR determinants; the carbapenemase gene bla_KPC_ and several other AMR genes were located on other plasmids. Concerningly, the ability for the virulence plasmids to be maintained in *K. pneumoniae* lineages suggests that once established in the MDR hospital outbreak-associated clones, they may become quite stable. The initial report of iuc+ KPC+ ST11 in China prompted multiple other groups to report detection of the same strain in their hospitals (59–61), suggesting this strain may indeed be emerging as a persistently hypervirulent and MDR form of *K. pneumoniae*. Genomic surveillance and control of the spread of such ‘dual-risk’ strains, or indeed even plasmids combining both characteristics of MDR and hypervirulence clearly needs to be reinforced; the present work will bolster efforts to understand and limit the emergence of infections caused by *K. pneumoniae* strains carrying the high virulence determinants aerobactin and salmochelin.

**List of Abbreviations**
MDR: multidrug-resistant
HA: healthcare-associated
CA: community-associated
AMR: antimicrobial resistance
MGEs: mobile genetic elements
HGT: horizontal gene transfer
ICEs: integrative and conjugative elements
ST: sequence type
CG: clonal group
MLST: multi-locus sequence typing
Ent: enterobactin
Ybt: yersiniabactin
Iuc: aerobactin
Iro: salmochelin
AbST: aerobactin sequence type
SmST: salmochelin sequence type
OR: odds ratio
CI: confidence interval

## DECLARATIONS

### Ethics approval and consent to participate

Not applicable.

### Consent for publication

Not applicable.

### Availability of data and material

All whole-genome sequences analysed in this study are publicly available in NCBI or the NCTC 3000 Project website (https://www.sanger.ac.uk/resources/downloads/bacteria/nctc/), accession numbers are listed in **Table S1**. Complete genome sequences generated for this study (summarised in **Table S3**) have been deposited in NCBI GenBank under accessions TBC. Accession numbers for the 12 reference plasmid sequences are listed in **Table S2**; the set of annotated sequences and the Mauve multiple alignment of these sequences are also deposited in FigShare (doi:10.6084/m9.figshare.6839981). The aerobactin and salmochelin MLST schemes are available in the *K. pneumoniae* BIGSdb database (http://bigsdb.pasteur.fr/klebsiella/klebsiella.html) and in the Kleborate distribution (https://github.com/katholt/Kleborate).

### Authors’ contributions

MMCL performed the majority of data analyses and wrote the paper together with KEH. RRW, KLW, SB, and KEH contributed additional data analysis, visualisation and interpretation. SB incorporated the novel MLST schemes into the BIGSdb. RRW and KEH wrote code. AJ contributed clinical isolates, data and interpretations. LMJ performed DNA extraction and nanopore sequencing. All authors edited and approved the paper.

## Acknowledgements

We thank the team of the curators of the Institut Pasteur MLST system (Paris, France) for importing novel alleles, profiles and/or isolates at http://bigsdb.pasteur.fr.

## Supplementary Tables

Table S1. Strain information for genomes included in this study.

Table S2. General features of reference plasmids or incomplete plasmid sequences carrying *iro* and/or *iuc*.

Table S3. Summary of replicon sequences from strains INF151, INF237 and INF078

Table S4. Aerobactin sequence types (AbSTs) and corresponding alleles.

Table S5. Salmochelin sequence types (SmSTs) and corresponding alleles.

Table S6. Representative Enterobacteriaceae genome sequences included in *iro* and *iuc* phylogenetic analysis

Table S7. Single nucleotide variants and nucleotide divergence (%) observed within (shaded in grey) and between the aerobactin-encoding *iuc* lineages.

Table S8. Single nucleotide variants and nucleotide divergence (%) observed within (shaded in grey) and between the salmochelin-encoding *iro* lineages.

Table S9. Summary of aerobactin-encoding iuc and salmochelin-encoding *iro* loci BLAST hit

Table S10. Prevalence of virulence loci and plasmid replication loci amongst strains with virulence plasmids

## Supplementary figures

**Figure S1.**
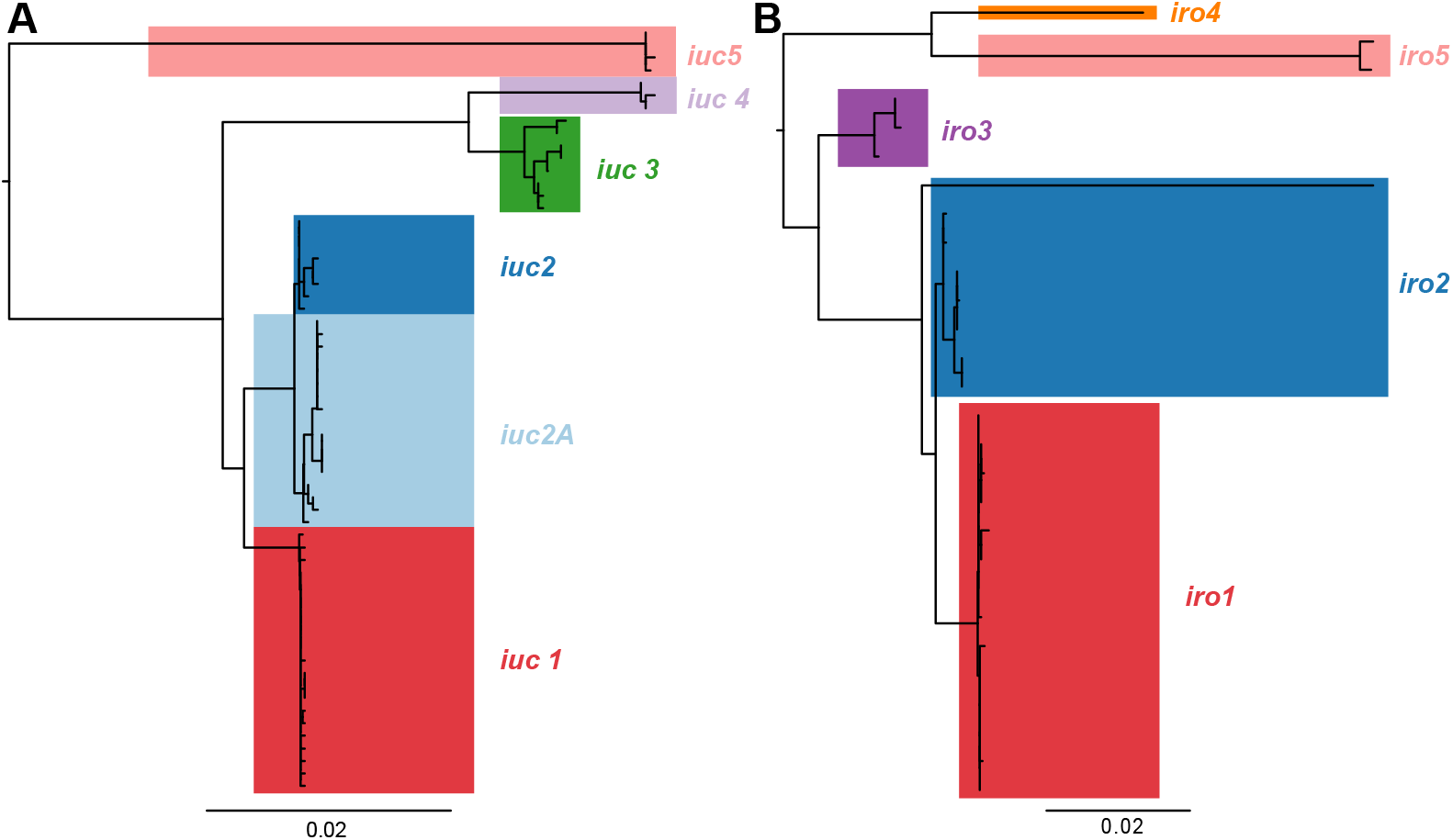
Phylogenetic relationships between the predicted amino acid sequences encoded by aerobactin (*iuc*) and salmochelin (*iro*) locus sequence types. Each tip represents a translated amino acid sequence for an aerobactin sequence type (AbST, in **a**) or salmochelin sequence type (SmST, in **b**). Lineages defined from nucleotide sequences (see tree in **Fig. 1**) are highlighted and labelled.

**Figure S2.**
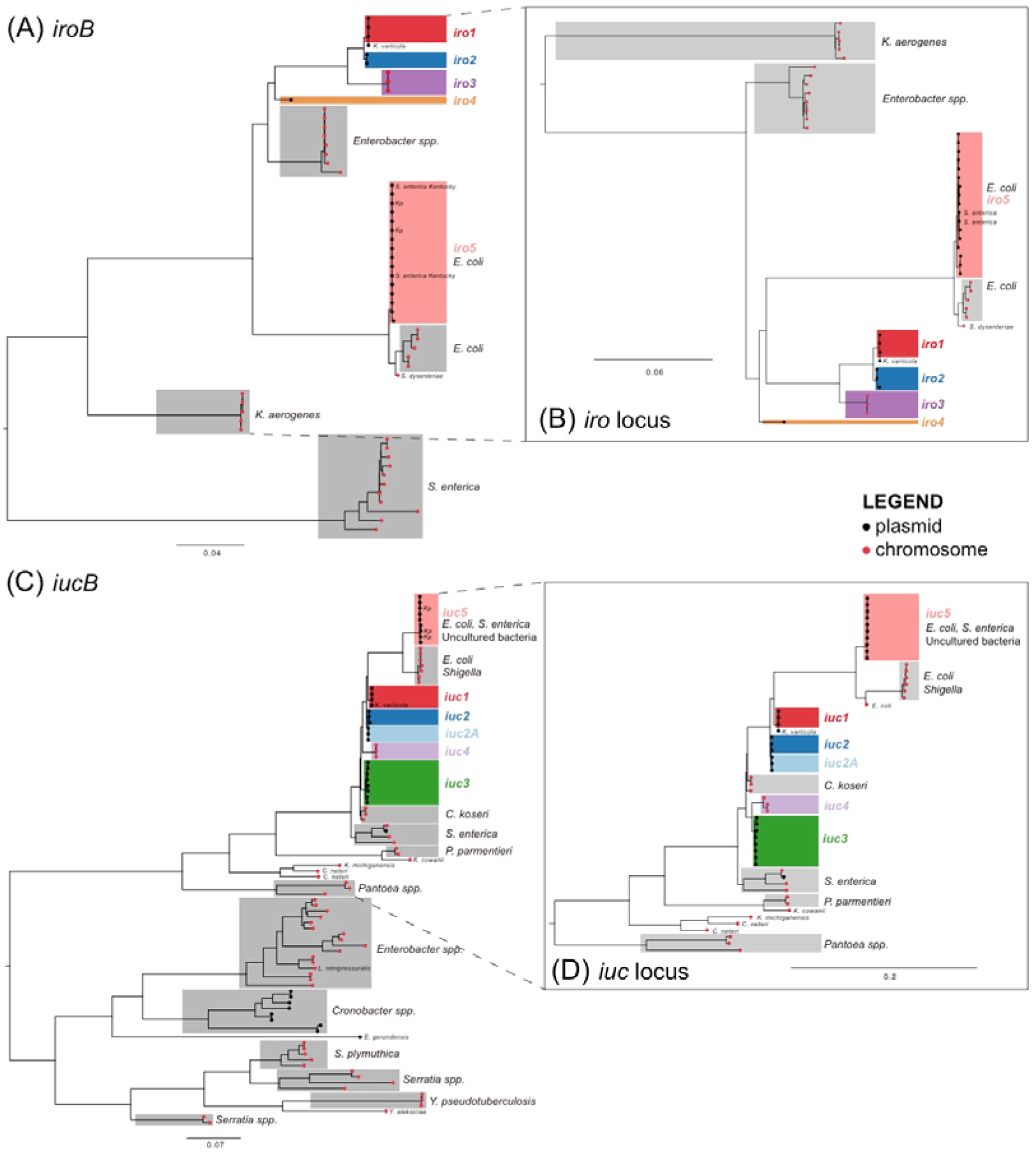
Phylogenetic trees for salmochelin and aerobactin encoding *iuc* locus in *K. pneumoniae* and other Enterobacteriaceae bacteria. Trees represent show a midpoint-rooted maximum likelihood phylogeny for representative sequences identified in various Enterobacteriaceae species (listed in **Table S6**). Tip colours indicate the genetic context of the locus: black=plasmid, red=chromosome. *K. pneumoniae iro* lineages defined in **Fig. 1** are coloured; other species-specific clades are highlighted in grey; individual labelled tips within highlighted clades indicate exceptions to the species label of the clade. Salmochelin trees were inferred using the *iroB* gene alone (panel **a**), which show a highly divergent form in *Salmonella*. Panel (**b**) Shows a tree inferred from all four genes of the typical *K. pneumoniae iro* locus (*iroBCDN*), excluding the distantly related *Salmonella* variant, to increase resolution within the group containing *Klebsiella*. Similarly, aerobactin trees were inferred using the *iucB* gene alone (panel **c**) to show the overall structure, and separately for the full set of genes in the *K. pneumoniae* locus (*iucABCD, iutA*) to provide greater resolution within the group containing *Klebsiella* (panel **d**).

**Figure S3.**
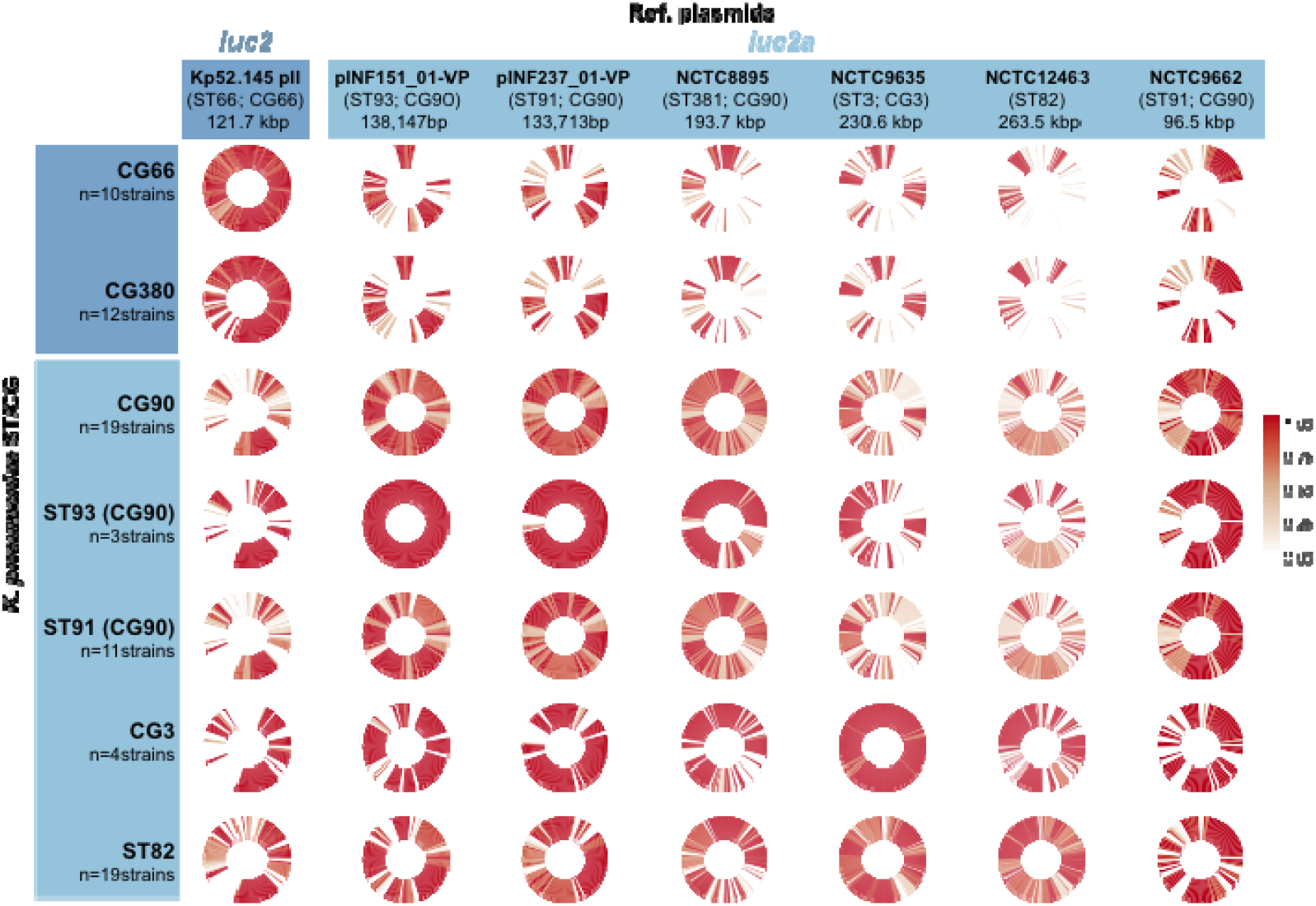
Conservation of coding sequences from KpVP-2 and *iuc2a+* reference plasmids amongst strains carrying plasmid-encoded *iuc2* or *iuc2a* loci. Cells show circularised heatmaps indicating the frequency of each gene in a given reference plasmid (column), amongst strains of a given chromosomal sequence type (ST) or clonal group (CG) (rows) that carry either *iuc2* (CG66, CG380) or *iuc2a* (others). Around each circle, genes are ordered by their order in the corresponding reference plasmid.

**Fig. S4.**
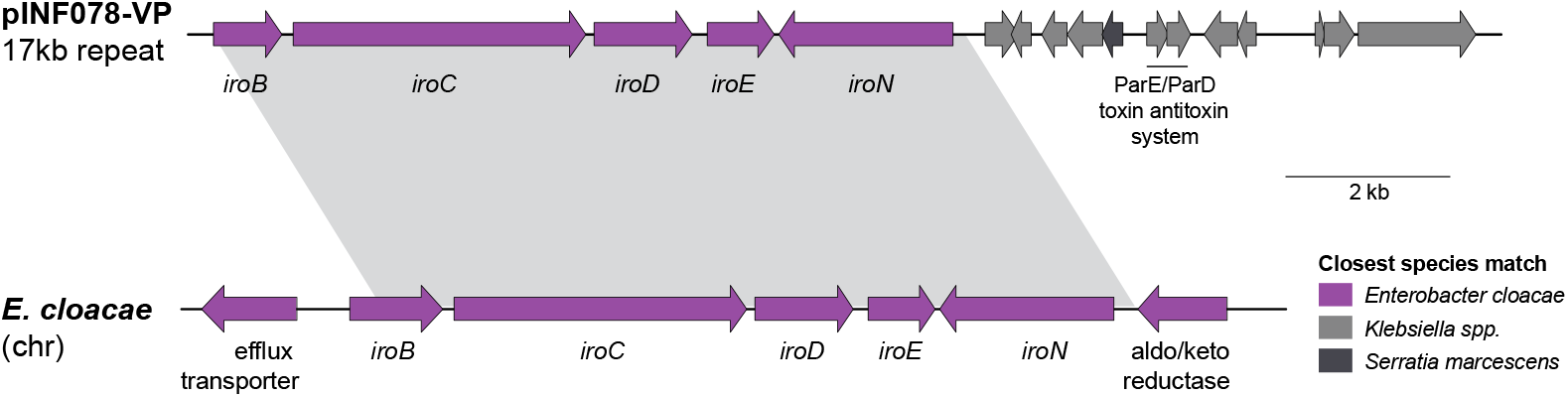
Genetic structure of 17 kbp repeat region in plasmid pINF078-VP and the chromosomally-encoded *E. cloacae iro* region. Shaded area indicates a homologous region of 95% nucleotide identity shared between the two sequences. Coding sequences are represented by the arrows and coloured according to the closest Enterobacteriaceae species match as indicated in the legend.

